# CDK1 and PLK1 co-ordinate the disassembly and re-assembly of the Nuclear Envelope in vertebrate mitosis

**DOI:** 10.1101/185660

**Authors:** Ines J de Castro, Raquel Sales Gil, Lorena Ligammari, Maria Laura Di Giacinto, Paola Vagnarelli

**Affiliations:** College of Health and Life Science, Research Institute for Environment Health and Society, Brunel University London, UB8 3PH, UK

## Abstract

Micronuclei (MN) arise from chromosomes or fragments that fail to be incorporated into the primary nucleus after cell division. These structures are a major source of genetic instability caused by DNA repair and replication defects coupled to aberrant Nuclear Envelope (NE). These problems ultimately lead to a spectrum of chromosome rearrangements called chromothripsis, a phenomenon that is a hallmark of several cancers. Despite its importance, the molecular mechanism at the origin of this instability is still not understood. Here we show that lagging chromatin, although it can efficiently assemble Lamin A/C, always fails to recruit Nuclear Pore Complexes (NPCs) proteins and that Polo-Like Kinase (PLK1) negatively regulates the NPC assembly. We also provide evidence for the requirement of PLK1 activity for the disassembly of NPCs, but not Lamina (A/C), at mitotic entry. Altogether this study reveals the existence of independent regulatory pathways for Lamin A/C and NPC reorganization during mitosis where Lamin A targeting to the chromatin is controlled by CDK1 activity (a clock-based model) while the NPC loading is also spatially monitored by PLK1.

## INTRODUCTION

During open mitosis the nuclear envelope breaks down at the end of prophase to allow the separation of sister chromatids and starts to re-assemble during anaphase (reviewed in [1]). Lamin A phosphorylation by cdc2/cyclinB [2, 3] allows lamina disassembly at the beginning of mitosis while multiple kinases appear to contribute to the disassembly of the NPC via phosphorylation of a crucial target, Nup98 [4]. During mitotic exit, NE reformation (NER) occurs as a sequential and multi-step process where NUPs assemble to the chromatin earlier than Lamin A [5] but the overall mechanism is still poorly understood (recently reviewd in [6] and [7]).

A few studies have shown that this mechanism is regulated by the concerted action of protein phosphatase 1 (PP1) [8] [9] and 2A (PP2A) [10]. This ordered assembly suggests that sequential dephosphorylation events must occur to allow macromolecular complexes to form and target them to the correct place at the right time. This could be achieved by activation of different phosphatases at different times during mitotic exit (clock model) or by the position of the segregating chromatin within the anaphase cell (spatial model) or a combination of both. However the molecular effectors are still not known. Moreover, recent work in Drosophila has shown that an Aurora B gradient appears to regulate a surveillance mechanism that prevents both chromosome decondensation and NER until effective separation of sister chromatids is achieved [11] and to delay Lamin A re-assembly [12].

Here we show for the first time that, in vertebrates, Polo-like kinase 1 (PLK1) is also participating in this NER control pathway, and that it is essential both for timely disassembly of the NPC at the onset of mitosis and for preventing the loading of the NPC on the chromatin of lagging chromosomes during mitotic exit.

This study proposes a sequential model for the NER where Lamin A targeting to the chromatin is under the control of the decreasing Cdc2/cyclinB activity (a clock-based model) while the NPC loading is also spatially regulated by PLK1 activity.

## RESULTS AND DISCUSSION

In order to understand the causes of MN instability, we have analysed the composition of the NE of MN in HeLa and U2OS cells by immunofluorescence.

While Lamin B1 was absent in all MN analysed in both cell lines (as previously reported [13]), we noticed heterogeneity among the MN in relation to the presence/absence of Lamin A and Importin βor the NPC marker mAb414, which recognizes several FG nucleoporins, with the majority of the MN lacking Importin β(Supplementary Figure 1 A, B). Although the absence of Lamin B1 could indicate that the MN have already been disrupted (as reported by Zhang et al [14]), the presence or absence of the other nuclear membrane (NM) components suggested that the loading of Lamin A could be uncoupled from that of nucleoporins.

Indeed, Lamin A and NPC seem to have different loading times.

In order to understand how this variability could arise, we have focused on anaphase cells containing lagging chromosomes or chromatin bridges, the most common cause for MN formation [15]. NER starts in anaphase in a sequential manner and the order of events have been well documented [5]. For example, NUPs (as judged by mAb414 staining) and Importin βrecruitment precedes Lamin A and it starts loading at the pole-ward side of the segregating chromatids as shown in Supplementary Figure 1 C, D.

Lagging chromatids arise from chromosome fragments without kinetochores or merotelic-attached chromosomes [16], dwelling in the middle of the anaphase spindle and causing a delay in division. Using several antibodies that recognize different NPC sub-complexes (m414 for FG containing NUPs, Nup93 for the inner ring complex, POM121 for the transmembrane NUPs and MEL28 for the Y complex [17], we have revealed that chromatin present in the middle of the anaphase/telophase spindle is competent for Lamin A assembly but fails to recruit either Importin βor NPC components (Figure 1 A-F). Therefore, while a fraction of interphase MN are loaded with NPC components (Supplementary Figure 1A-B), lagging chromatin is not (Figure 1 A-F) but still does assemble Lamin A indicating that the normal order of assembly (first NPC then Lamin A) is not maintained in this type of chromatin. These observations suggest that: 1) the process of Lamina re-assembly and NPC reassembly after mitosis are independent of each other and they are under different regulatory systems; 2) recruitment of NPC components to the anaphase chromatin is negatively regulated directly or indirectly by factors (possibly kinases) acting at the central spindle or limited by the small amount of chromatin present. We discarded the latter hypothesis since we could identify cells with a substantial mass of lagging chromatin which was still negative for NPC components (Figure 2 A).

**Figure 1:**
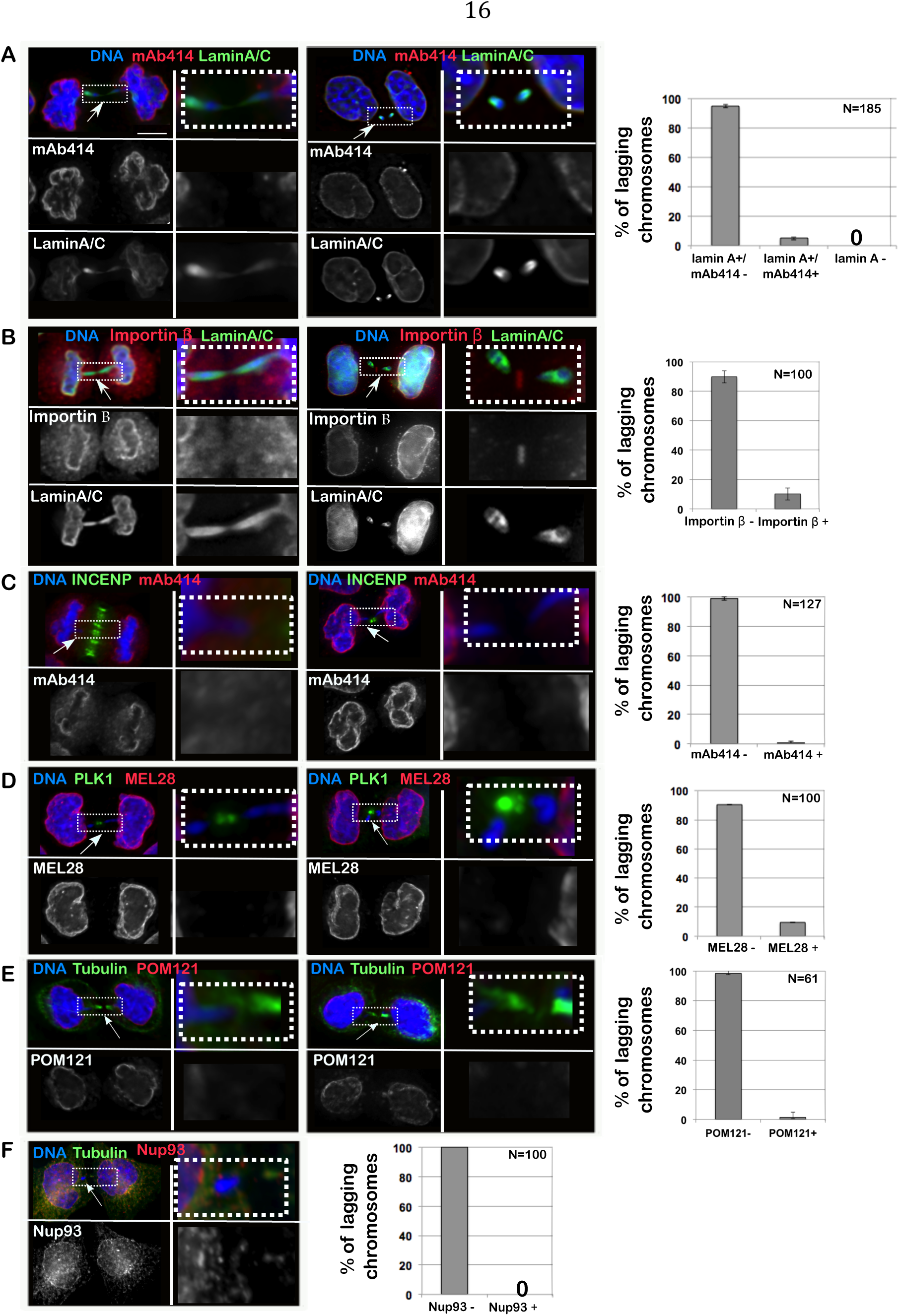
Importin β and NPC do not assemble on lagging chromatin in anaphase. HeLa cells were fixed and stained for Lamin A/C, Importin β and several components of the NPC and the presence of the staining on chromatin bridges was recorded. (A-F) HeLa cells in telophase/cytokinesis showing chromatin bridges and stained with: Lamin A/C (green) and mAb414 (red) (A); Lamin A/C (green) and Importin β (red) (B); INCENP (green) and mAb414 (red) (C); PLK1 (green) and MEL28 (red) (D); GFP:POM121 (red) and α-tubulin (green) (E); α-tubulin (green) and NUP93 (red) (F); scale bar 10μm. The insets indicate the region of the picture shown at higher magnification in the adjacent column. A‘-F’: quantification of the experiment in (A). The graphs show the distribution of the different classes as a mean of 3 independent experiments; the error bars represent SD and the number indicates the total cells counted.

**Figure 2:**
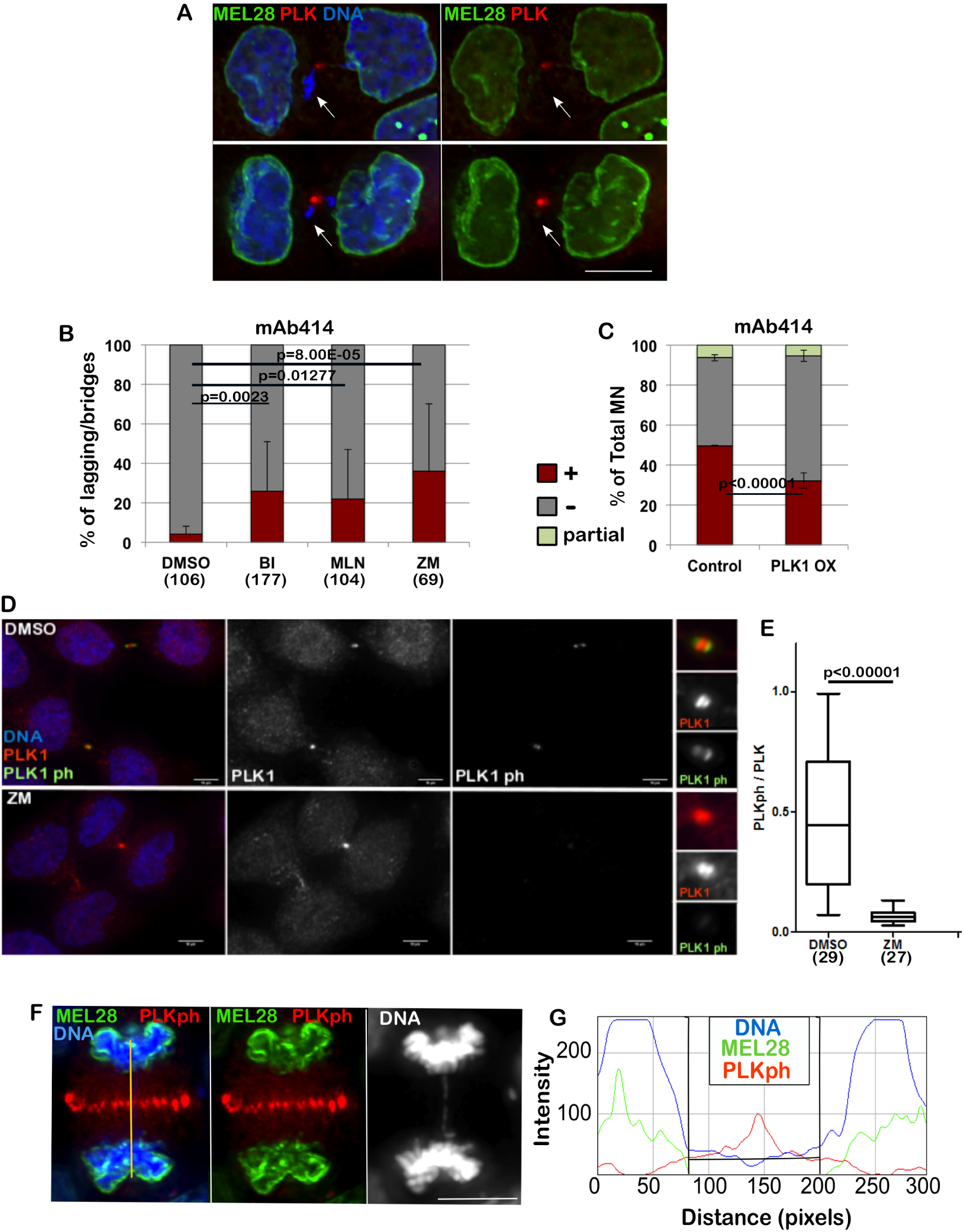
PLK1 and Aurora B regulate the loading of NPC. A: HeLa cells were fixed and stained for MEL-28 and PLK1. The arrows indicate the lagging chromatin and no presence of the NPC component. Scale bar 10μm. B: HeLa cells were treated with DMSO, BI 2536 (BI), MLN0905 (MLN), both targeting PLK1, or ZM447439 (ZM), targeting Aurora B, for 4h then fixed and stained for mAb414. The lagging chromatin in telophase cells was analysed for presence or absence of the staining. The error bars indicate the SD between 3 replicates and Fisher exact test was used for statistical analyses. C: HeLa cells were transiently transfected with mCherry:PLK1 (PLK1 OX) and fixed and stained for mAb414 24h post-transfection. The graph shows the percentage of total micronuclei (MN) positive (brown) or negative (grey) for the staining. The error bars indicate the SD between 3 replicates and Fisher exact test was used for the statistical analyses. D: HeLa cells were treated with either DMSO or ZM447439 (ZM) for 30 min then fixed and stained for PLK1 (red) or PLK1ph (green). On right: zoom in of the staining at the midbody. Cells at the midbody stage were used for the analyses. Scale bar: 10 μm. E: quantification of the experiment in (D). The signal intensity ratio between phosphorylated PLK1 and total PLK1 at the midbody in control (DMSO) and Aurora B inhibited (ZM) cells. Mann-Whitney U test was used for statistical analyses. F: HeLa cells were stained for MEL28 (green) and PLK1ph (red). Anaphase cells showing absence of MEL28 staining on the lagging chromatin. The yellow line indicates the region used for the intensity profile in (G). G: Intensity profile of the staining in (F) showing accumulation of MELS28 only when PLK1ph decreases.

We therefore started investigating a possible role for kinases as negative regulators of the NER. In Drosophila, it has recently been shown that Aurora B acts in anaphase to delay Lamin A and B re-assembly when chromatin is trapped in the midzone [11] [12]. In contrast, our experiments indicate that Lamin A can successfully assemble on lagging chromatin (Figure 1B). We therefore tested if inhibition of different kinases present in the anaphase midzone/midbody could allow NPC loading on lagging chromatin in vertebrates. Inhibition of PLK1 or Aurora B was able to release the inhibitory effect and allowed part of the NPC loading to the lagging chromatin (Figure 2B). While inhibition of Aurora B could have been expected based on the experiments conducted in Drosophila, the effect caused by PLK1 inhibition is surprising since this kinase has never been reported to act on NE reformation nor has it been shown to be able to generate a gradient in anaphase. Supporting our experiment, PLK1 was shown to be recruited to DNA tethers of acentric chromatin during cell division [18]. A further support to the implication of PLK1 in NPC regulation was provided by experiments where by overexpression of RFP:PLK1 in HeLa cells we were able to see a significant reduction of NPC at the NE of MN (Figure 2 C). This observation could either suggest that both kinases regulate a subset of NUPs or that one kinase activity depends on the other.

While it is known that Aurora B activates PLK1 at kinetochores in early mitosis both in Drosophila and human cells [19], its regulation in late mitosis is poorly understood. Only recently it has been suggested that in Drosophila, Map205 sequesters Polo kinase on central spindle microtubules and that upon phosphorylation mediated by Aurora B, active Polo is recruited to the midbody where it targets substrates locally. This model has the potential to serve as the regulatory mechanism for PLK1 and NPC loading. However, the vertebrate orthologue of Map205 is not known. We have therefore tested if a crosstalk between Aurora B and PLK1 could also exist in human cells. To this purpose we have treated HeLa cells for 30 min with the Aurora B inhibitor ZM447439 (ZM) and quantified the level of Phospho-PLK1 at the midbody relative to total PLK1. The results clearly show that Aurora B inhibition causes a significant decrease in Phospho-PLK1 at the midbody (Figure 2 D, E) thus indicating that, as in Drosophila, activation of PLK1 in the midbody depends on Aurora B. These results could potentially indicate that the phosphorylation status of PLK1 could affect NPC re-assembly. In fact, NPC deposition around the chromatin appears to occur away from the presence of PLK1ph, where PLK1ph levels are below a threshold (Figure 2 F, G).

In order to test the relevance of protein kinases in envelope re-assembly further, we analysed the process in a system where spatial clues are removed. Cells blocked in mitosis can be induced to biochemically recapitulate several physiological steps of mitotic exit when cyclin-dependent kinase 1(CDK1) is inactivated ([9, 20, 21]. To this purpose, we used a chicken DT40 cell line engineered to express an analog sensitive version of CDK1 [22] thus removing the spatial clues from the system. This allele (CDK1^as^) can be specifically inhibited by the bulky ATP analogue 3MB-PP1. The cell line was also transfected with GFP:Lamin A (DT40 does not express Lamin A), blocked in prometaphase with nocodazole and the accumulation of NPC and lamina at chromatin was checked. These cells were then treated either with DMSO, BI2536, 3MB-PP1, ZM447439 or BI2536 + 3M-PP1 for 15 minutes and fixed and stained with mAb414 or Importin β. In DMSO treated cells, both Lamin A and the NPC or Importin βare dispersed in the cytoplasm (Figure 3 A-D). Upon inhibition of CDK1, Lamin A is re-localised onto the chromatin but no further accumulation is produced by co-inhibition of PLK1 (Figure 3D, E). Inhibition of CDK1 also triggers NPC recruitment on the chromatin in this system but the additional inhibition of PLK1 produces a nuclear-rim like and more substantial accumulation of mAb414 and Importin βto the chromosomes (Figure 3 A-C). No effect can be observed by inhibiting either PLK1 alone or Aurora B alone (Figure 3 A-C). Inhibition of the kinases was effective within the 15 min treatments as judged by a parallel experiment where cells were treated with single drugs and MG132; the mitotic cells were negative for H3S10ph or showed monopolar spindles upon treatment with ZM447439 or BI2536 respectively (Supplementary Figure 2A) as expected when Aurora B or PLK1 are inactive. This data suggests that CDK1 is solely responsible for maintaining Lamin A off the chromosomes in mitosis but additional phosphorylations by PLK1 or kinases downstream of PLK1 are required to prevent the NPC loading in late mitosis. Taken together the observations so far provided, we can conclude that a conserved mechanism sustained by PLK1 activity (localised at the midzone in normal mitotic exit) negatively regulates the NPC reformation during anaphase. However this mechanism does not prevent Lamin A recruitment indicating therefore that the two processes are independent and they are monitored by different kinases/phosphatases.

**Figure 3:**
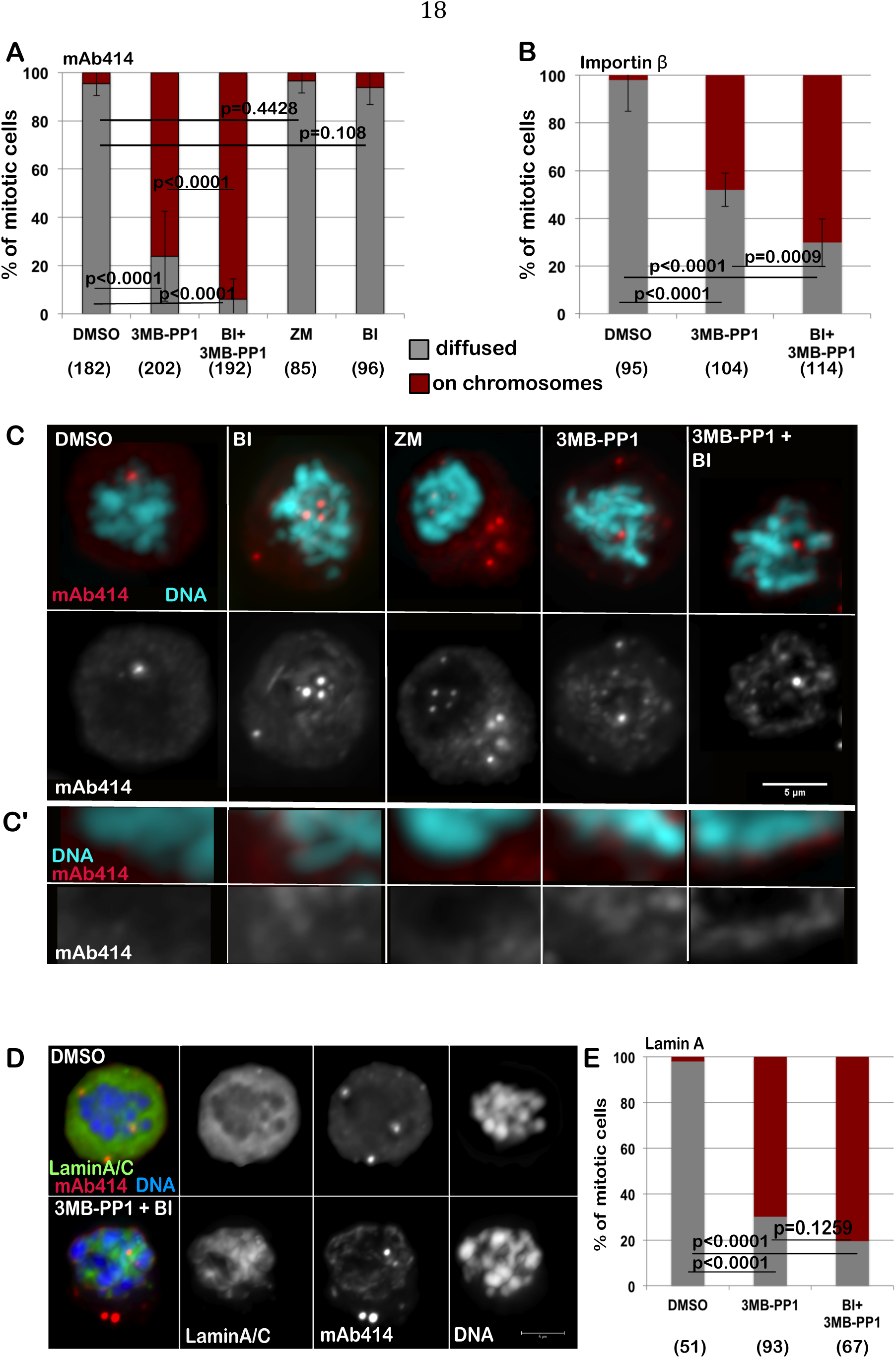
Inhibition of CDK1 and PLK1 in prometaphase cells are both required for the complete loading of the NPC and Importin β on the chromatin. A-B: DT40CDK1^AS^ were arrested in nocodazole for 4h then treated for 15 min with DMSO, 3MB-PP1, ZM447439 (ZM), BI2536 (BI) or 3MB-PP1+BI2536 (BI+3MB-PP1). The cells were then stained either for mAb414 (A) or Importin β (B) and the localization of the staining was recorded. The error bars indicate the SD between 3 replicates and Fisher exact test was used for the statistical analyses. C: representative images of the experiment shown in (A). mAb414 (red) and DNA (blue). Scale bar 5 μm. C’: Enlargements of the pictures in (C). D: DT40CDK1^AS^ were transiently transfected with GFP:LaminA and treated as in A. The localization of GFP:LaminA was recorded. The panels show representative images of cells treated only with DMSO (upper panels) or 3MB-PP1 + BI (bottom panels). E: quantification of the experiment in (D). The error bars indicate the SD between 3 replicates and Fisher exact test was used for the statistical analyses.

This data reconciles with what is known for Lamin A dynamics in mitosis where CDK1 has been found responsible for phosphorylation of lamins, a prerequisite for NE disassembly. At mitotic entry, the phosphorylation of lamins at specific sites by CDK1 and protein kinase C (PKC) is required to drive disassembly of the lamina [23, 24] [3] [2] and dephosphorylation of the CDK1 mitotic sites by protein phosphatase 1 (PP1) is required for lamina assembly during the telophase/early G1 transition [25]. It is therefore expected that just inhibition of CDK1 would license Lamin A deposition on the chromatin.

Given the fact that PLK1 could play a role in NPC re-organisation during mitotic exit, we hypothesized that it could also be important for the dissolution of NPC at mitotic entry. Although a requirement for PLK1 in mitotic entry has been suggested, with p150^Glued^ being one of the substrates involved in this process [26], little is known about PLK1 involvement in the dissolution of the NPC at the onset of mitosis. Nup98, a nucleoporin phosphorylated by multiple kinases, is essential for Nuclear Envelope Breakdown (NEBD) and is also a PLK1 substrate; however, mutation of the two identified PLK1 sites had no impact on NEBD in vitro [4]. Nevertheless, we have noticed that mcherry:PLK1 accumulates at the nuclear periphery in prophase, at a time consistent with its involvement in NE re-organisation (Figure 3 A, B). A similar observation was reported in Drosophila, where Polo kinase also localises to the NE at mitotic entry [19].

To test therefore if PLK1 was required for the NPC dissolution at mitotic entry, we inhibited CDK1 with 3MB-PP1 in the DT40CDK1^AS^ cell line for 6h to block cells at the G2/M boundary. After 30 min release, to allow the cells enter mitosis, we inhibited PLK1 or Aurora B for 15 min and analysed the distribution of Importin βor mAb414 in mitotic cells (Figure 4 C). While in normal and Aurora B inhibited prometaphase/metaphase cells mAb414 and Importin βare both diffused and on the spindle, in PLK1 inhibited cells Importin βand the NUPs are either still around the condensed chromosomes or in distinct patches within the cytoplasm (Figure 4 D-G). Inhibition of Aurora B does not produce any detectable effect on NPC dispersal (Figure 4 E, F). However, neither inhibition of PLK1 nor Aurora B affects dispersal of Lamin A (Figure 4 J-L). The same results were obtained in HeLa cells. In this system we have inhibited either PLK1 (using either BI2536 or MLN0905) or Aurora B for 1 h or 2 h and then stained the cells for mAb414 and Lamin A/C. Again the inhibition of only PLK1 causes a delay in the dissolution of the NPC but does not affect the disassembly of Lamin A/C (Figure 4 H-L). These results clearly indicate that also at mitotic entry different kinases are required to complete NEBD and that the breakdown of Lamin A and the NPC follow different pathways.

**Figure 4:**
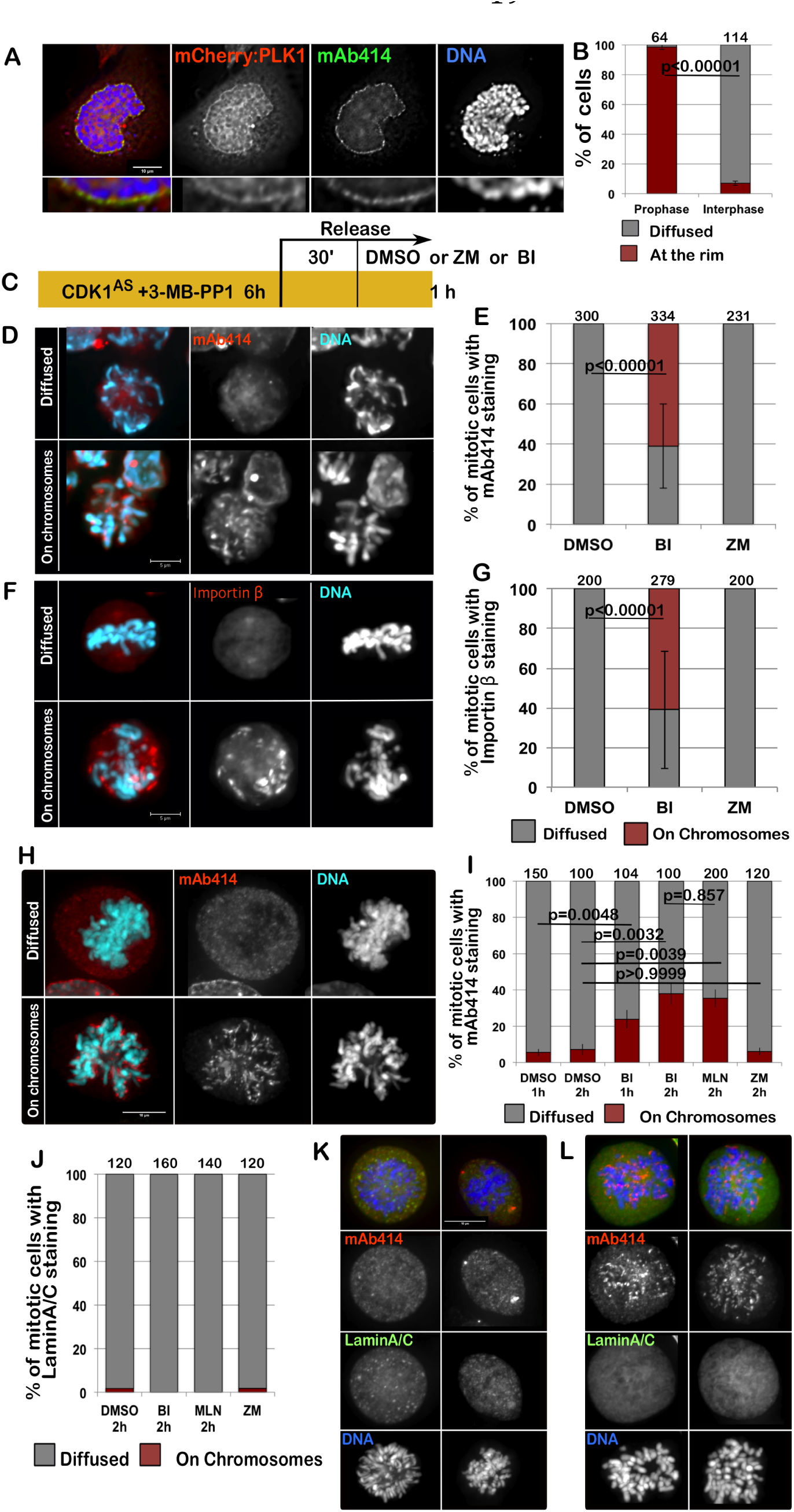
PLK1 is required for the complete disassemble of the NPC at the onset ofs mitosis. A: HeLa cells were transfected with mCherry:PLK1 (red) then fixed and stained with mAb414 (green). The nuclear localization of PLK1 was then recorded in interphase and prophase cells. The panels show representative images of a prophase cell. The insets below are magnification of the image above. B: quantification of the experiment in (A); Fisher exact test was used for the statistical analyses. C: scheme of the experiment. D-G: DT40CDK1^AS^ after the treatments as in (C) were fixed and stained for Ab414 or Importin β. Inhibition of PLK1 compromises the dispersal of the NPC in mitotic cells. The panels show representative images for the staining with mAb414 (D) or Importin β (F) in DMSO (diffused) or BI (on chromosomes)–treated cells (D); E and G: quantification of the experiments in D and F. The error bars indicate the SD between 3 replicates and Fisher exact test was used for the statistical analyses. H-I: HeLa cells were treated with either DMSO, BI 2536 (BI), MLN0905 (MLN) or ZM447439 (ZM) for 1 or 2h then the cells were fixed and stained for mAb414 (red). The distribution of mAb414 classified as “diffused” or “on chromosomes” was quantified (I). Representative images of a “diffuse” (upper panels) or “on chromosome” (lower panel) pattern are presented in (H). J-L: HeLa cells were treated with either DMSO, BI 2536 (BI), MLN0905 (MLN) or ZM447439 (ZM) for 1 or 2h then the cells were fixed and stained for mAb414 (red) and Lamin A/C (green). The distribution of Lamin A classified as “diffused” or “on chromosomes” was quantified (J). The error bars indicate the SD between 3 replicates and Fisher exact test was used for the statistical analyses. Representative images of DMSO (K) or BI (L)–treated cells are shown.

**Figure 5.**
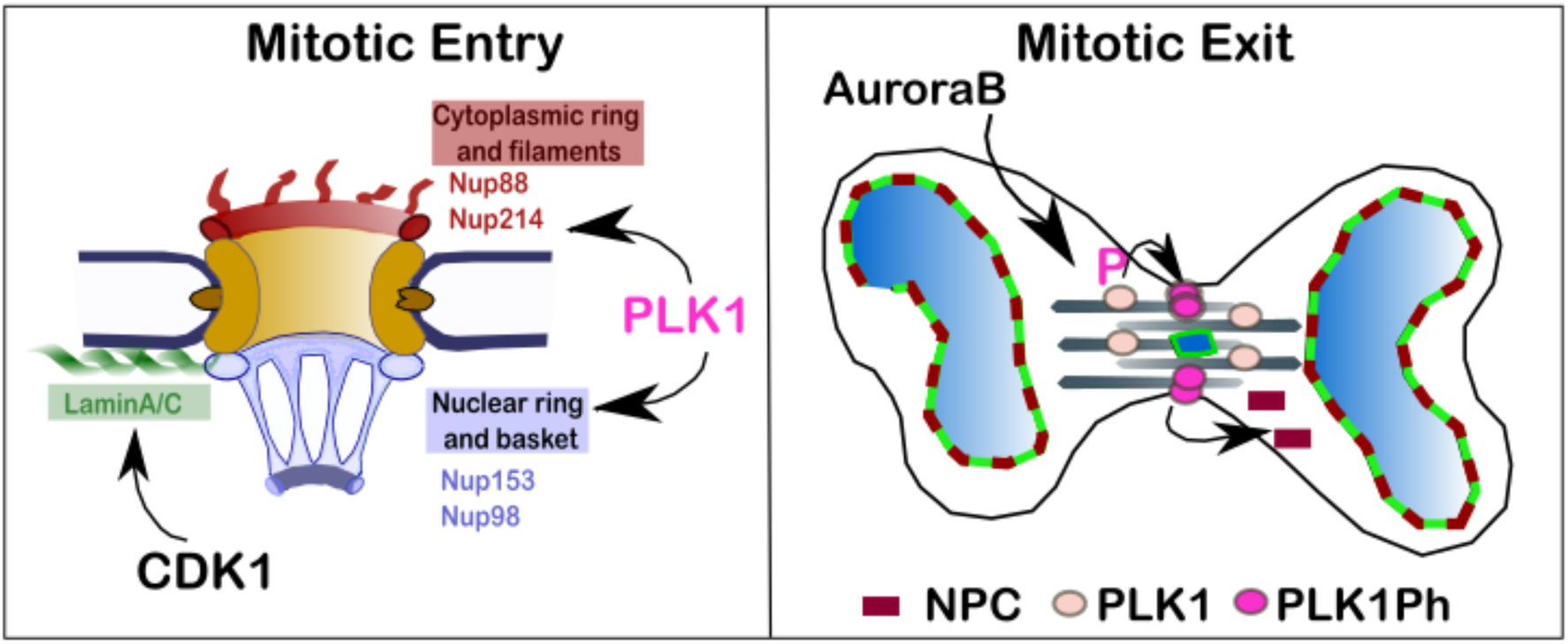
Proposed model for the regulation of NPC re-organisation in mitosis. At mitotic entry PLK1 localises at the nuclear rim in prophase where phosphorylates a subsetofnucleoporinscontributingtothedisassemblyoftheNPC.CDK1 phosphorylates Lamin A and is important for the NE disassembly. At mitotic exit, CDK1 inhibition is essential for triggering NE re-assembly and is sufficient for targeting Lamin A/C to the chromatin in vertebrates. The complete NPC re-assembly requires in addition the inhibition of PLK1 which activity is sustained by Aurora B at the cleavage furrow.

Here we have shown that Lamina A/C and NPC dissolution and reformation are controlled by different mechanisms. CDK1 is necessary for Lamin A breakdown and its inhibition is sufficient for targeting it onto the chromosomes. However, the NPCs require the additional phosphorylation by PLK1 for their full dispersal at onset of mitosis, and PLK1 inactivation for the re-assembly at mitotic exit. These observations are consistent with the fact that mitotic phosphorylation of several nucleoporins depends onPLK1as suggested by published proteomic studies [27-29] and supported by previous observations linking lack of PLK1 to delays of NEBD in human cells [30], C. elegans [31] [32] and in mouse oocytes maturation [33]. However this is the first demonstration that NPC and Lamin A/C are behaving in an independent manner. At mitotic exit the NPC re-assembly is spatially controlled by the presence of active PLK1 whereas Lamin A is regulated via a “clock” mechanism based on inactivation of CDK1 without any topological control; in fact it is not important where the chromatin is within the anaphase cell to be competent for Lamin A binding. Although the PLK1-non-phosphorylable mutants of Nup98 did not show an effect in inhibiting NEBD in vitro [4], it is possible that several phosphorylation events on different nucleoporins have additive effects and altogether contribute to the full NPC disassembly. Therefore, mutations in single components might not be sufficient to produce major defects.

This study therefore provides mechanistic insights into the pathway on NE reorganisation in vertebrates and the identification of the specific counteracting phosphatases will be an important aspect to pursue in order to understand the full picture. It is possible that PLK1 negatively regulates the activity of phosphatases as well. Hints towards this direction are that some NPC components have been shown to be substrates for PP2A-B55 [34] but the sites identified are not the ones sensitive to PLK1 inhibition [29].

It is worth noting that Repo-Man/PP1, which activity is regulated by Aurora B [35], can bind to Nup153 [36] and is important for Importin βrecruitment on the lagging chromatin [9]; this complex could also represent a potential phosphatase candidate. Importantly, our findings have also major implications in genome stability. As we have shown, the MN that originate from merotelic attached chromosomes are inevitably going to have a compromised nuclear barrier from the start as their import/export machinery is impaired. These MN will then be a major source of genomic instability as elegantly demonstrated recently by the Pellman Lab [14]. These findings could therefore add extra value to the use of PLK1 inhibitors for the treatment of specific cancer types arising from MN formation.

## ACKNOWLEDGMENTS

We would like to thanks Mar Carmena (Edinburgh), Eric Shirmer (Edinburgh) and Joanna M Bridger (Brunel, London) for critical reading of the manuscript.

MLDG was supported by a Placement ERASMUS fellowship.

The Vagnarelli Lab is funded by BBSRC (grant BB/K017632/1 to PV).

---

**Supplementary Figure 1.**
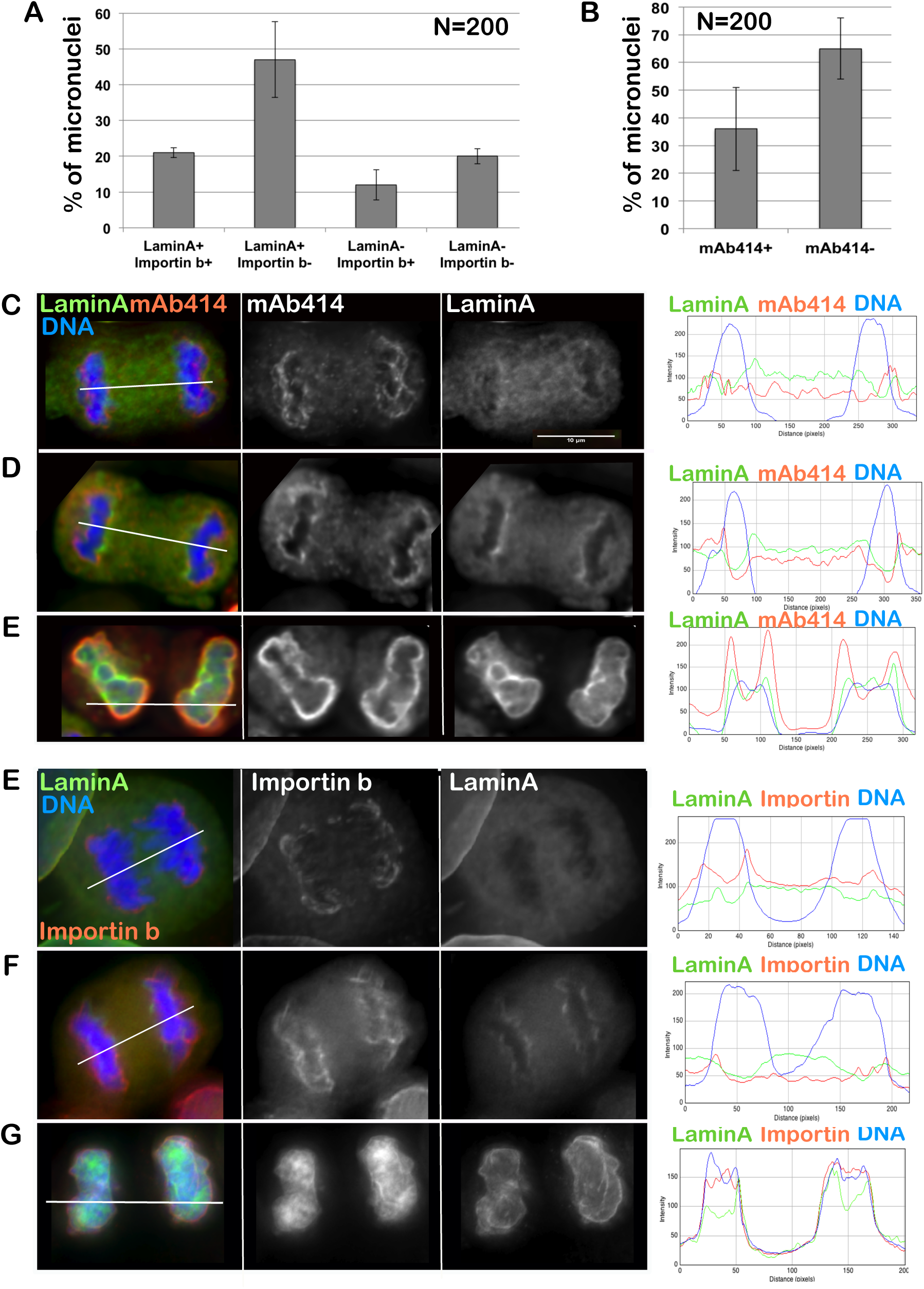
Importin β and Nups are loaded before Lamin A/C on the anaphase chromosomes starting from the pole-ward side of the chromatin. A-B: HeLa cells were stained for Importin β, mAb414 and Lamin A/C. The presence/absence of the staining in MN was recorded and plotted. The error bars indicate the SD between 3 replicates. C-H: HeLa cells were stained for mAb414 (C-E) or Importin β (E-G) and Lamin A/C. Intensity profiles on the distribution of the markers along the two sister chromatids. In early anaphase mAb414 (C) and Importin β (F) accumulate around the chromosomes starting from the side toward the pole but no signal is present on the telomeric side (the middle of the cell) of the chromosomes; at this stage the Lamina is still diffused; later in anaphase mAb414 (D) and Importin β (G) have completely surrounded the chromatin and the lamina is loading. In cytokinesis both NM components are fully assembled onto the chromosomes (E and H).

**Supplementary Figure 2.**
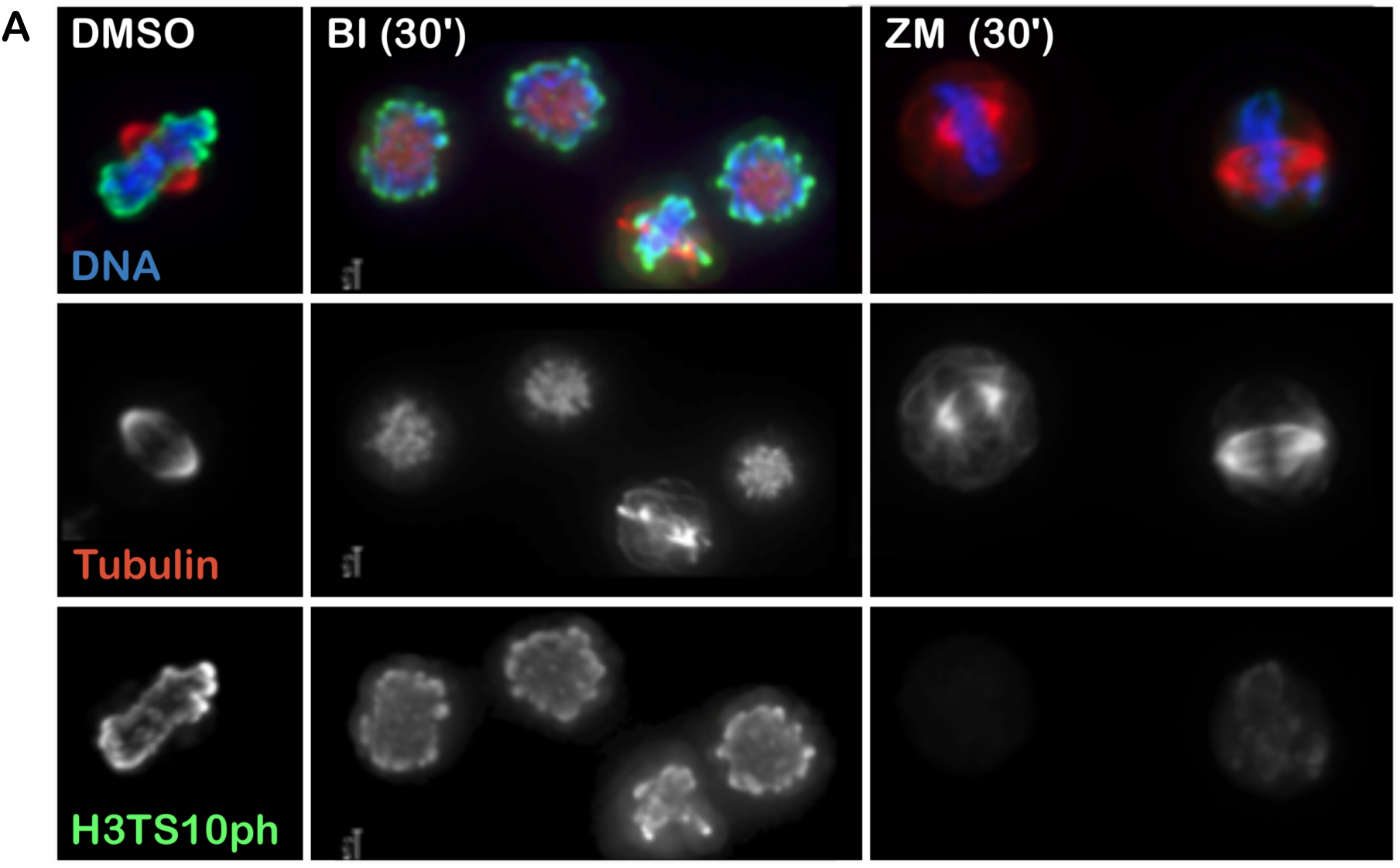
Short inhibition of PLK and AuroraB. DT40CDK1AS cells were blocked with 3MB-PP1 for 6 h then released for 60’ in MG132 containing medium. After BI or ZM were added for 15’. The cells were then fixed and stained with anti αtubulin and anti H3S10ph antibodies.

## MATERIALS AND METHODS

**Cell Culture**

HeLa cells were cultured in DMEM supplemented with 10% FBS and DT40 cells were cultured in RPMI1640 supplemented with 10% FBS and 1% chicken serum at 37°C and 39°C respectively in 5%CO_2_. DT40 cells DT40CDK1^AS^ cells were provided by Dr Helfrid Hochegger (Sussex, UK). HeLa GFP:POM121 cell line was kindly provided by Kathrine Ullamn (Utha, USA).

Transient transfections for DT40 were conducted as previously described [9]. For synchronization, DT40 cells DT40CDK1^AS^ cells were treated with 100 mM of 3MB-PP1 (Millipore) for 4h or with 100ng/ml nocodazole (Sigma).

BI2536 (Stratech), ZM447439 and Y-27632 (Calbiochem), MLN0905 (Sellckechem) were used at 100nM, 2μM, 40μM, and 35μM respectively for the time indicated in each experiment.

mCherry-PLK1-N-16 was a gift from Michael Davidson (Addgene plasmid 55119)

**Immunofluorescence and microscopy**

For immunofluorescence, cells were fixed in 4% PFA and processed as previously described [9]; antibody incubations were carried out in 1% BSA-phosphate-buffered saline (PBS) with 1%BSA and 0.01% TWEEN for 1 h at 37°C.

Fluorescence-labelledsecondaryantibodieswereappliedat1:200(Jackson ImmunoResearch).

The primary antibodies were used as follows:

Anti Nup153 (mouse QE5, Abcam, ab24700) 1:300; mAb414 (mouse Covance 1:500, MMS-120R); Anti-Importin β (monoclonal 3E9, Abcam, ab2811) 1:1000; anti-Lamin A/C (rabbit monoclonal, Abcam ab108595); anti-Ser10ph (rabbit, Upstate Biotechnoloy,#06-570) 1:200; anti-αtubulin (monoclonal B512, Sigma, T5168) 1:1000; anti-PLK1 (Abcam, ab175808) 1:1000; anti-PLK1ph (Abcam, ab39068) 1:300; anti-INCENP (gift from WC Earnshaw, Edinburgh) 1:500; Mel28 (Gift from M. Platani, Edinburgh) 1:500; Nup93 (Santa Cruz, GC-292099) 1:100.

3D data sets were acquired on a wide-field microscope (NIKON Ti-E super research Live Cell imaging system) with a NA 1.45 Plan Apochromat lens. The data sets were deconvolved with NIS-Element AR software. 3D data sets were converted to Quick Projections or presented as single plane, exported as TIFF files, and imported into Adobe Photoshop for final presentation.

For quantification of the staining (Figure 2 C-D), an area containing the midbody was select and the total staining in the red and blue channel was measured. Another set of measurements was obtained from the same areas in a region outside the cell (background) and subtracted by the intensities in the midbody.

The graphs were generated in Excel or R-studio, exported as png and imported in Inkscape for figure presentation.

